# Preferential consumption of benthic cyanobacterial mats by Caribbean parrotfishes

**DOI:** 10.1101/2022.11.09.515834

**Authors:** Joshua C. Manning, Sophie J. McCoy

## Abstract

Cyanobacteria are ubiquitous on coral reefs and perform important ecosystem functions. Benthic cyanobacterial mats (BCMs) have become increasingly abundant on degraded reefs. While epilithic and endolithic benthic cyanobacteria are the primary trophic resource for many parrotfishes, mat-forming benthic cyanobacteria are generally considered unpalatable to reef fishes. Regardless, recent studies have documented substantial grazing of BCMs by reef fishes, including parrotfishes. Here, we observed foraging in five Caribbean parrotfishes on the fringing coral reefs of Bonaire, Netherlands, to investigate BCM consumption relative to other benthic substrates. All species overwhelmingly targeted reef substrates composed of algal turfs and crustose coralline algae, which are typically associated with epilithic and endolithic microalgal and cyanobacterial communities. Additionally, three species preferentially consumed BCMs. As such, our work is consistent with and provides direct evidence supporting the recently proposed trophic role for parrotfishes as microphages. Contrasting observations of reef fishes avoiding substrates dominated by BCMs on other reefs suggests variation in the palatability of BCMs to grazing reef fishes, or species-specific differences in preference for these potentially nutritional trophic resources.

## Introduction

Cyanobacteria are a ubiquitous component of benthic coral reef communities and serve many important functional roles, including in nitrogen fixation, carbonate cycling, and primary productivity (Charpy et al. 2012). As coral reefs have become degraded by rising ocean temperatures and localized stressors (e.g., eutrophication), mat-forming benthic cyanobacteria have increased in abundance on many coral reefs (de Bakker et al. 2017; Reverter et al. 2020, 2022). Benthic cyanobacterial mats (BCMs) overgrow and smother benthic organisms, including corals (Puyana and Prato 2013). Overgrowth of corals causes significant tissue damage and reduces growth rates (Titlyanov et al. 2007). BCMs may also act as reservoirs for potentially pathogenic bacteria (Cissell et al. 2022), and the increased cover of BCMs has been correlated with an increase in coral disease (Reverter et al. 2020). Additionally, benthic cyanobacteria inhibit coral larval recruitment and reduce their likelihood of survival (Kuffner et al. 2006; Ritson-Williams et al. 2020). As such, there is growing interest and urgency in understanding the dynamics of mat formation and persistence.

Many mat-forming benthic cyanobacteria (e.g., *Lyngbya* spp.) produce secondary metabolites that deter grazing by generalist herbivores (Thacker et al. 1997; Nagle and Paul 1998, 1999). However, mat forming benthic cyanobacteria are also a potentially rich source of nutrients for herbivores due in to lower C : N content than many other algal resources (Atkinson and Smith 1983; Capper et al. 2006), and experimental work confirms that fish will consume them in equal abundances to algal resources when secondary metabolites are undetectable (Capper et al. 2006). Production of secondary metabolites in mat-forming benthic cyanobacteria varies spatiotemporally (Nagle and Paul 1999; Paul et al. 2007), even at small scales (Capper et al. 2006). Thus, the composition of mat-forming bacterial communities and variation in the production of secondary metabolites by these communities may drive geographic differences in the prevalence and consumption of cyanobacterial mats (Cissell and McCoy 2022).

Microscopic photoautotrophs, including benthic cyanobacteria, are the primary nutritional target of parrotfishes (Clements et al. 2016; Nicholson and Clements 2020, 2021). A recent study on the fringing coral reefs of Bonaire, Netherlands, found that BCMs comprised a substantial proportion of bites taken by striped parrotfish (*Scarus iseri*), and were frequently consumed by other fishes, including blue parrotfish (*Scarus coeruleus*; Cissell et al. 2019). Frequent consumption of BCMs suggests that they may be an important and relatively novel nutritional resource for parrotfishes, particularly in Bonaire, where their cover has steadily increased in recent decades (de Bakker et al. 2017). However, BCMs have been found to disrupt natural grazing processes on other reefs where they are rarely consumed (Ford et al. 2021; Ribeiro et al. 2022). As such, additional research is needed to determine the importance of BCMs as a resource for parrotfishes.

Here, we observed foraging behavior in five Caribbean parrotfishes common to the fringing coral reefs of Bonaire: *Scarus vetula, Sc. taeniopterus, Sc. iseri, Sparisoma viride*, and *Sp. aurofrenatum*. We investigated the relative importance of BCMs to these five parrotfish species compared to other components of their diets, including both terminal phase (TP) and initial phase (IP) fish for three of those species (*Sc. vetula, Sc. taeniopterus*, and *Sp. viride*). We discuss our findings in light of recent studies on the nutritional ecology of parrotfishes.

## Materials and Methods

### Study location and site characteristics

We collected data from five fringing coral reef sites along the leeward coast of Bonaire during May – July 2019: Bachelor’s Beach, The Lake, Angel City, Aquarius, and Invisibles (Fig. 1). We estimated the cover of different foraging substrates at each site from 1/16 m^2^ photoquadrats placed at 1 m intervals along 10 m transect lines that were haphazardly placed and run perpendicular to the reef slope at ~10 m depth [Supplemental Information (SI): Table S1]. Photoquadrats were not moved to artificially select for hard substrates (e.g., Steneck et al. 2019) and, therefore, included sediment and rubble habitat where BCMs are often observed. We randomly allocated 49 points to each photoquadrat and identified the benthic substrate (i.e., macroalgae, coral, etc.) under each point using the software Coral Point Count with Excel Extensions (Kohler and Gill 2006).

**Fig. 1:**
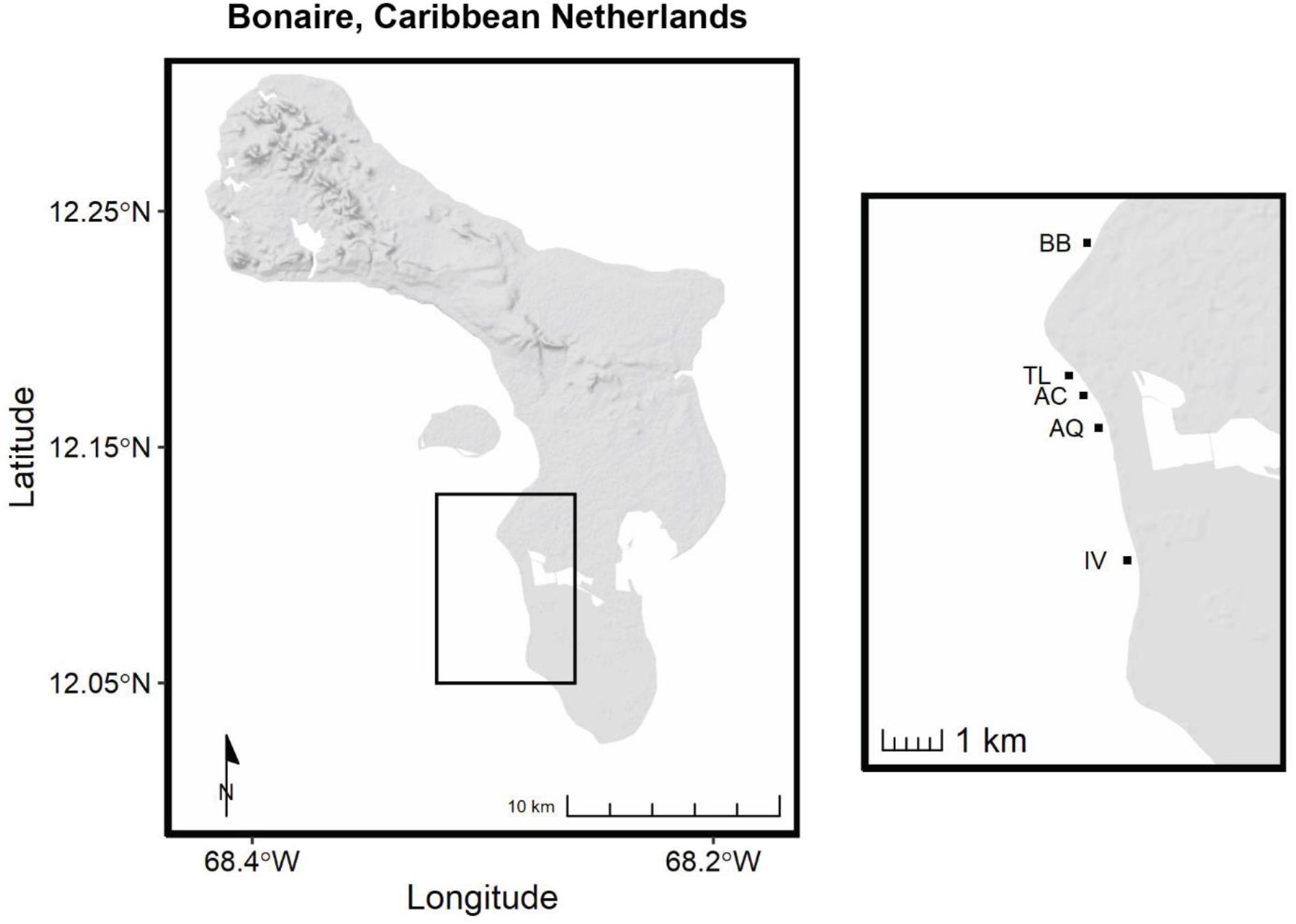
A map of our field sites in Bonaire, Caribbean Netherlands: Angel City (AC), Aquarius (AQ), Bachelor’s Beach (BB), Invisibles (IV), and The Lake (TL).

We estimated the density of parrotfishes at ~10 m depth at each site by conducting visual censuses along eight 100 m^2^ band transects (*sensu* Steneck et al. 2019). We swam each transect at a constant rate (~5 m min^-1^) and counted all TP and IP parrotfishes greater than 5 cm fork length, placing them into size class bins (6-10, 11-20, 21-30, and 31-40 cm). We used the densities and the mean fork length (8, 15.5, 25.5, and 35.5 cm) of each bin to estimate biomass of each species using published length-weight relationships (Bohnsack and Harper 1988). Mean (± SE) parrotfish densities (counts 100 m^-2^) and biomass (g 100 m^-2^) are presented for each site in the SI (Table S2).

### Foraging observations

We conducted video-recorded, behavioral observations of five Caribbean parrotfish species: *Sc. taeniopterus, Sc. vetula, Sp. viride, Sc. iseri*, and *Sp. aurofrenatum*. Behavioral observations were made during peak parrotfish foraging periods (1000 – 1600 hrs; Bruggemann et al. 1994b, 1994a). We observed 128 territorial TP parrotfishes, including *Sc. taeniopterus, Sc. vetula*, and *Sp. viride* at all five study sites, and *Sc. iseri* and *Sp. aurofrenatum* at two study sites (Aquarius and Invisibles). Territorial TPs forage in fixed diurnal home ranges, from which intraspecific TPs are largely excluded (Manning and McCoy *in review*). In contrast, non-territorial (transient) TPs are often chased along the reef by territory holders, making observation difficult and likely influencing foraging behavior. Thus, we excluded transient fishes from our analyses. To explore the effect of ontogenetic phase on foraging behavior, we also observed 34 large IP *Sc. vetula*, *Sc. taeniopterus*, and *Sp. viride* at two sites (Aquarius and Invisibles). The total number of behavioral observations and the mean (± SE) duration of these observations are reported for each species and ontogenetic phase in the SI (Table S3).

Focal parrotfishes (TP or IP) were identified haphazardly at ~10 m depth on SCUBA at each site and allowed to acclimate to diver presence for ~ 1-2 mins, during which time the observer estimated its standard length to the nearest cm (Table S3). We then followed the focal fish for 12.9 ± 0.2 mins (mean ± SE observation time, n = 162) from ~2 m away and recorded foraging behavior with a GoPro Hero 4 Silver (GoPro, Inc; 4k resolution) attached to a ‘selfie-stick’, to be analyzed later in the behavioral analysis software BORIS (v. 7.9.8; Friard and Gamba 2016). All bites were counted and the identity of the substrate targeted during each bite was recorded.

Substrates targeted were grouped as follows: BCM, algal turfs and crustose coralline algae, fleshy erect macroalgae (hereafter, macroalgae), live coral (Scleractinia and Milleporidae), sediment, soft coral (Gorgonia), sponge, and other (all other bites on benthic substrates). BCMs have been defined as macroscopic, cohesive colonies growing over sediment and hard reef substrates (including live benthic organisms), which are taxonomically composed primarily of Cyanobacteria and Proteobacteria (Cissell and McCoy 2021). Algal turfs are heterogenous communities that include filamentous turf algae and cyanobacteria, small non-calcified crusts, macroalgal propagules, and associated detritus (Bruggemann et al. 1994c; Wilson and Bellwood 1997; Adey 1998; Fricke et al. 2011). Algal turfs often, but not always, contained sediments. In a few rare occurrences, fishes were observed cropping algal turf from the surface of sponges without any obvious removal of sponge tissue. These bites were scored as algal turf. Bites scored as sediment had no obvious epilithic algal/cyanobacterial filaments within or BCMs atop them. When identification of a bite target was impossible or questionable because the view of the bitten substrate was obscured by another structure or by the body of the focal fish, the substrate target was scored as “unknown”. Parrotfishes frequently consumed the feces of planktivorous reef fishes (i.e., coprophagy). We include these bites in our analyses of foraging rates, but we have discussed their importance elsewhere (Manning and McCoy 2022).

### Statistical analyses

We analyzed all foraging data in R (v. 4.0.2; R Core Team 2020). We investigated the effect of site, species, and ontogenetic phase on bite counts with an additive generalized linear model fit to a negative binomial distribution with the log of the observation time included as an offset (glmmTMB v. 1.0.2.1; Brooks et al. 2017). We also fit a reduced model including only the species for which we had observations of both ontogenetic phases. The reduced model included site, species, ontogenetic phase, and a species by ontogenetic phase interaction as fixed effects. We checked both models for overdispersion and zero-inflation (DHARMa v. 0.3.3.0; Hartig 2020) and for multicollinearity (performance v. 0.6.1; Lüdecke et al. 2020). We then tested for the significance of the fixed effects in each model using Type III Wald’s χ^2^ tests (car v. 3.0-10; Fox and Weisberg 2019). We chose not to include body size as a predictor in our models because we focused our sampling focused on larger individuals of the two ontogenetic phases rather than a representative range of body sizes (Fig. S1). However, we discuss the potential importance of body size in explaining foraging differences within and among species.

We calculated Chesson’s α electivity index for each individual (dietr v. 1.0; Borstein 2019) to investigate foraging preferences based on the relative abundance of the different foraging targets at each site (Chesson 1983). Chesson’s α electivity indices were calculated only for the eight benthic substrates targeted by parrotfishes during foraging observations (bites on unknown substrates and fecal matter in the water column were excluded from this analysis). A Chesson’s α electivity index of 1/8 (i.e., 1/number of categories) represented no preference. Mean Chesson’s α electivity indices (± 95% CI) were plotted by species and ontogenetic phase to visualize preference or avoidance for targeted substrates. Finally, we quantified the mean proportion of bites taken on BCMs for each species and the number of foraging bouts with 5 or more consecutive bites on BCMs to determine whether parrotfishes were sampling BCMs rather that selectively foraging upon them.

## Results

*Sc. vetula*, *Sc. taeniopterus*, *Sc. iseri*, *Sp. viride*, and *Sp. aurofrenatum* were the only five parrotfish species observed at all of our study sites during surveys of fish abundance and account for the majority (> 96%) of the parrotfish biomass at these sites (Table S2). Foraging rates differed significantly by species and ontogenetic phase (full model: χ^2^ = 803.79, df = 4, p < 0.001 and χ^2^ = 40.39, df = 1, p < 0.001, respectively). Foraging rates of *Scarus* spp. were higher than foraging rates of *Sparisoma* spp. (Fig. 2) and there was a significant interactive effect of species and ontogenetic phase when foraging rates were compared among *Sc. vetula*, *Sc. taeniopterus*, and *Sp. viride* (reduced model: χ^2^ = 8.24, df = 2, p = 0.02). Foraging rates were higher for IP than for TP *Sc. vetula* and *Sc. taeniopterus*, but there was no effect of ontogenetic phase on foraging rates in *Sp. viride* (Fig. 2). There were no differences in total foraging rates among sites.

**Fig. 2:**
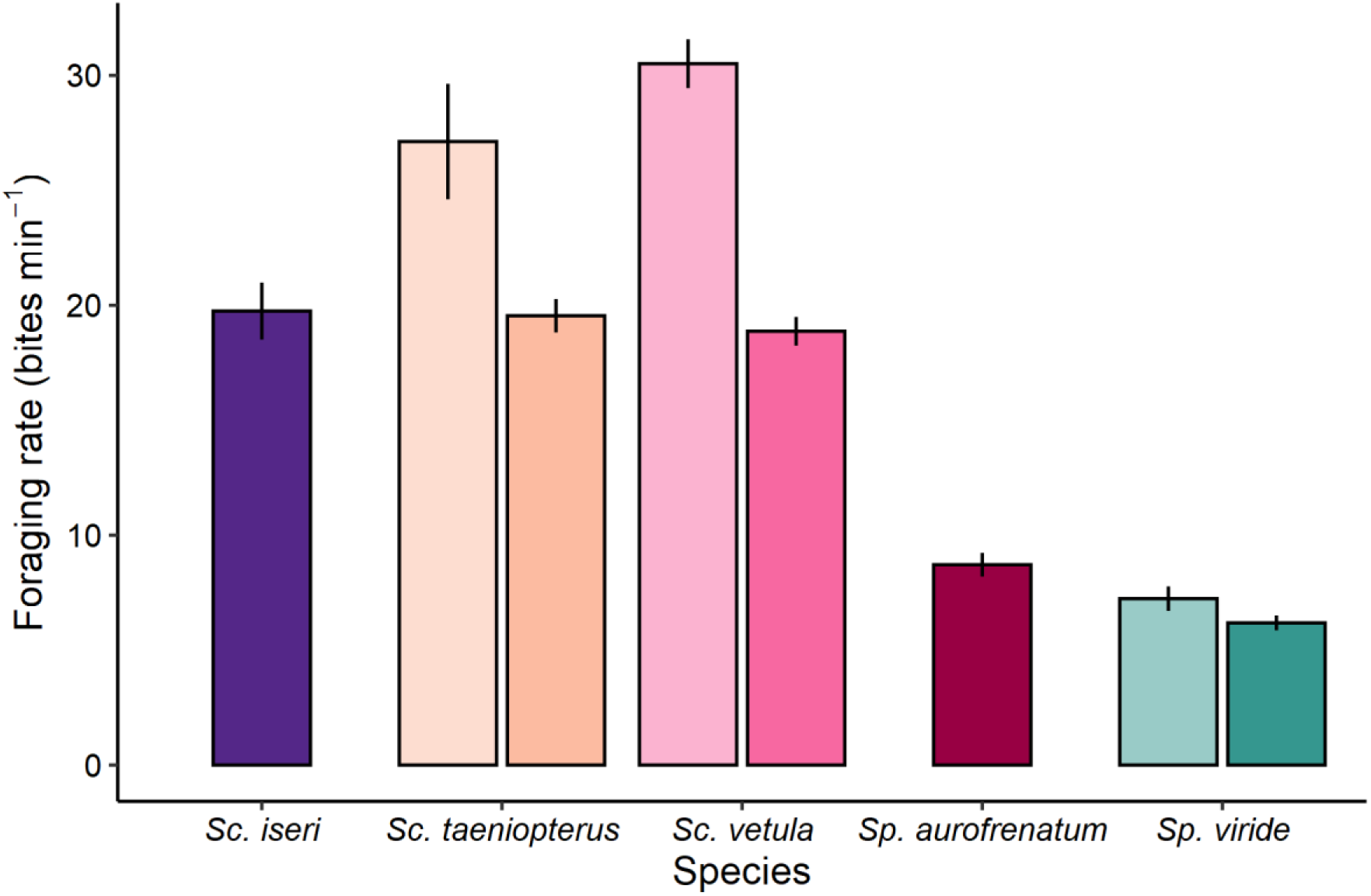
Mean (± SE) foraging rates (bites min^-1^) for TP and IP (when observed) *Sc. taeniopterus, Sc. vetula, Sc. iseri, Sp. aurofrenatum*, and *Sp. viride*.

We observed all five study species consuming BCMs, primarily from the sediment, but also from atop hard substrates (Fig. 3). BCMs were a preferred foraging substrate for TP and IP *Sc. iseri*, *Sc. taeniopterus*, and *Sc. vetula* (Fig. 4), despite their relatively low coverage on the reef (< 9%; Table S1). Bites on BCMs were between 7.1 ± 2.3 and 13.1 ± 1.4 % (mean ± SE, *Sc. iseri* and *Sc. taeniopterus*, respectively) of the total bites taken on benthic substrates for these three species (Fig. 3). BCMs comprised only 1.5 ± 0.3 and 6.0 ± 1.6 % of the bites taken by *Sp. viride* and *Sp. aurofrenatum*, respectively, and were not a preferred foraging substrate for either of these species (Figs. 3 and 4). *Scarus* spp., particularly *Sc. taeniopterus* and *Sc. vetula*, had many more feeding bouts with 5 or more consecutive bites on BCMs than either *Sparisoma* spp. (Fig. S2).

**Fig. 3:**
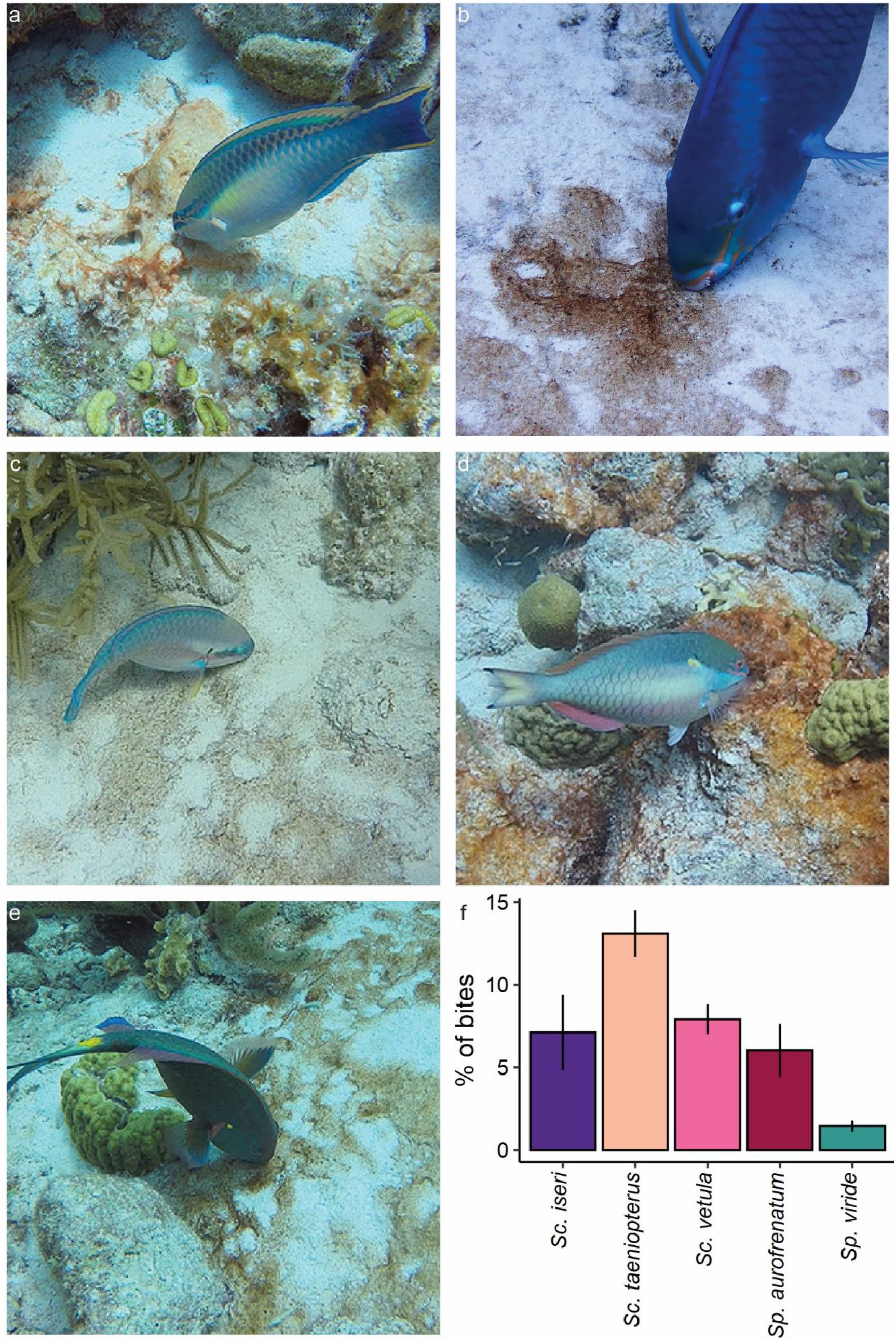
Examples of TP *Sc. taeniopterus* (a), *Sc. vetula* (b), *Sc. iseri* (c), *Sp. aurofrenatum* (d), and *Sp. viride* (e) taking bites of BCMs on the fringing coral reefs of Bonaire, and (f) the mean (± SE) percentage of bites taken on benthic cyanobacterial mats by each species.

**Fig. 4:**
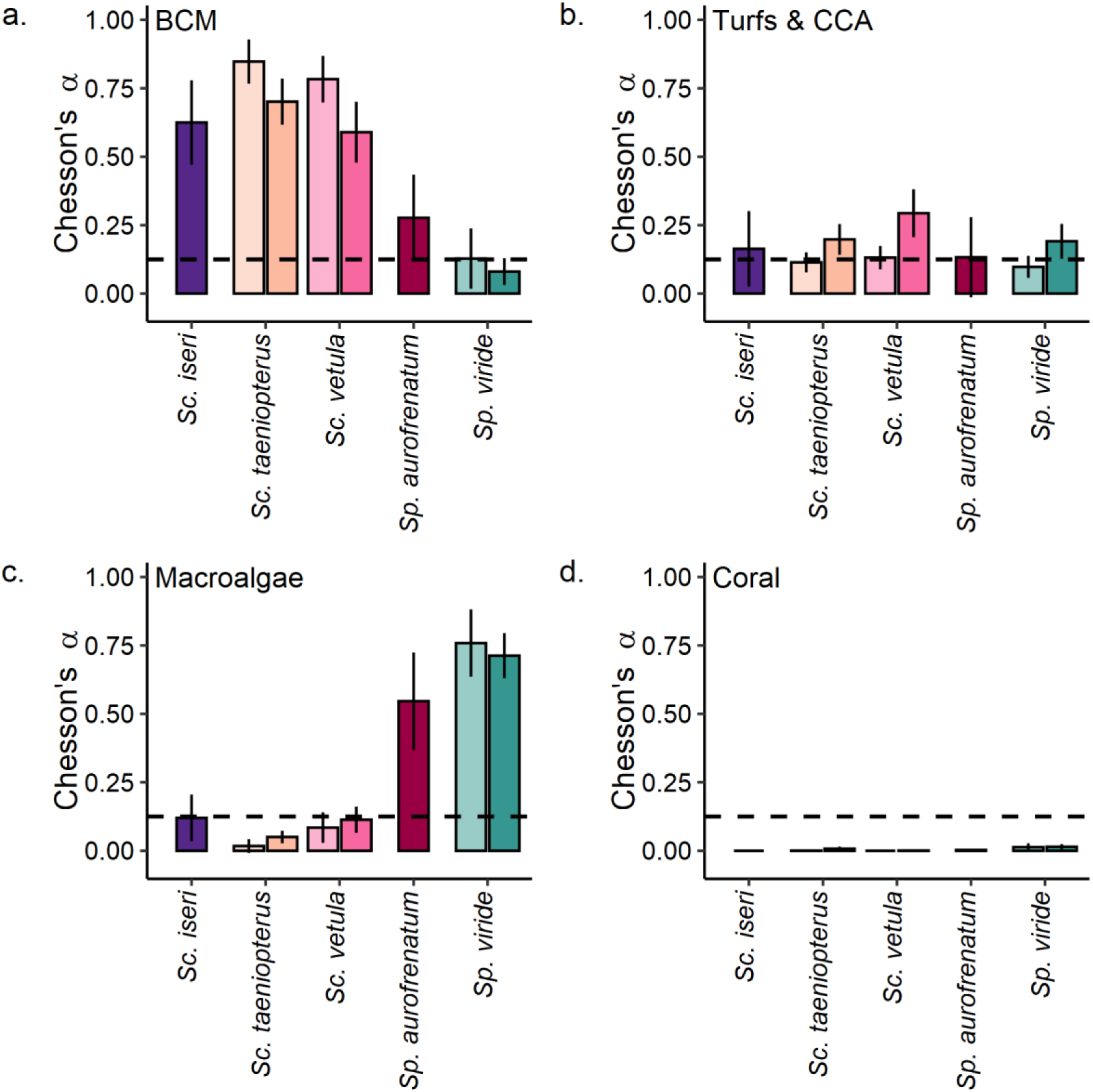
Mean (± 95% CI) Chesson’s α electivity index for terminal phase (TP) and initial phase (IP, when observed) *Sc. taeniopterus*, *Sc. vetula*, *Sc. iseri*, *Sp. aurofrenatum*, and *Sp. viride* and four major substrates: (a) BCMs, (b) algal turfs and crustose coralline algae, (c) macroalgae, and (d) live coral. The dashed line represents no preference (1/n; n = 8 foraging substrates).

The majority of the bites taken by all five species targeted substrates characterized as algal turfs and crustose coralline algae (69.5 ± 7.7 to 91.1 ± 0.9 %, mean ± SE; *Sc. iseri* and *Sc. vetula*, respectively). Algal turfs and crustose corallines were largely consumed in accordance with their abundance on the reef, though there was some evidence for preferential targeting by TP *Sc. taeniopterus* and TP *Sc. vetula* (Fig. 4). Both *Sp. viride* (TP and IP) and *Sp. aurofrenatum* preferentially consumed macroalgae (Fig. 4), which accounted for 7.9 ± 1.0 and 8.5 ± 2.1% of their bites, respectively. In contrast, *Sc. taeniopterus* (TP and IP) avoided macroalgae, while *Sc. vetula* (TP and IP) and *Sc. iseri* showed no preference or avoidance (Fig. 4). All species avoided live coral substrates (Fig. 4), though 3.3 ± 0.7 % of the bites taken by *Sp. viride* were on live corals.

## Discussion

As a disturbance that creates bare space, parrotfish grazing maintains reef substrates in cropped early successional states. This is considered critical for maintaining healthy and resilient coral reefs and facilitating coral larval recruitment (Mumby et al. 2006; Adam et al. 2015a). However, foraging rates and preferences vary both within and among parrotfish species (Bruggemann et al. 1994b; Bonaldo et al. 2006; Afeworki et al. 2013; Adam et al. 2015b; Smith et al. 2018). Here, we found species-specific differences in parrotfish foraging rates. Specifically, *Scarus* spp. foraged more frequently than *Sparisoma* spp. Ontogenetic phase and body size are also known to affect foraging rates in parrotfishes (Bruggemann et al. 1994a, 1994b; Afeworki et al. 2013), and there is some evidence that electivity for foraging substrates may differ by ontogenetic phase (Smith et al. 2018). For 3 of our study species (*Sc. taeniopterus, Sc. vetula*,and *Sp. viride*), we explored the effect of ontogenetic phase on foraging patterns. Overall, we found that foraging rates for *Sc. vetula* and *Sc. taeniopterus* were greater in initial phase than in terminal phase individuals, and our data suggested an inverse relationship between body size and total foraging rates, consistent with prior work (Bruggemann et al. 1994a, 1994b; Bonaldo et al. 2006; Afeworki et al. 2013). We found little evidence of ontogenetic differences in foraging preferences. So, we have chosen to focus our discussion on species differences in foraging preferences.

The abundance of BCMs has increased globally on coral reefs, including in Bonaire (de Bakker et al. 2017; Ford et al. 2018; Reverter et al. 2022). Mat-forming benthic cyanobacteria are known to produce secondary metabolites and have traditionally been considered unpalatable to reef fishes (Nagle and Paul 1998, 1999). As such, reef fishes have not been expected to play an important role in controlling mat proliferation, despite their importance in controlling reef macroalgal abundances. Ford et al. (2021) found that the presence of BCMs significantly reduced foraging by herbivorous reef fishes. In contrast, Cissell et al. (2019) documented extensive foraging on BCMs by multiple reef fishes, including by multiple parrotfishes in Bonaire. Consistent with Cissell et al. (2019), we have provided evidence that three common Caribbean parrotfishes in Bonaire preferentially consume BCMs growing over sediment and hard reef substrates.

Conflicting evidence of BCM consumption may reflect geographic differences in the composition of these mat communities or the production of secondary metabolites within them. Few studies have investigated the taxonomic composition of BCMs (but see Biessy et al. 2021; Cissell and McCoy 2021), and much of the research surrounding the chemical defenses of BCMs has focused on secondary metabolites isolated from a few mat-forming species (e.g., *Lyngbya* spp.; Thacker et al. 1997; Nagle and Paul 1999). The production of secondary metabolites can also vary, even at small spatiotemporal scales (Nagle and Paul 1999; Capper et al. 2006; Paul et al. 2007). Thacker et al. (1997) hypothesized that fishes may sample BCMs to effectively determine the concentrations of deterrent chemicals without over-ingesting toxins. When secondary metabolites are undetectable, reef fishes consume BCMs in equal abundance to less defended algal resources (Capper et al. 2006). In this study, we recorded multiple foraging bouts during which focal fishes, particularly *Scarus* spp., took many consecutive bites (> 5) of BCMs in Bonaire. Therefore, it appears that these fishes are selectively targeting BCMs rather than sampling them, suggesting that BCMs at the sites studied here are not deterring foraging chemically or that the parrotfishes are not affected by the chemical defenses present. These findings provide direct evidence in support of the hypothesis that parrotfishes are microphages that target protein rich microscopic photoautotrophs, primarily cyanobacteria (Clements et al. 2016).

Observational studies have often identified epilithic turf algae and fleshy macroalgae as the primary foraging substrates for many parrotfishes (Bruggemann et al. 1994c, 1994b; Adam et al. 2015b). In this study, parrotfishes overwhelmingly targeted turf algal communities and crustose coralline algae. Benthic cyanobacteria are a major component of epilithic (‘turf’) and endolithic algal communities on the reef benthos, where they are important primary producers (Adey 1998; Diaz-Pulido and McCook 2002; Tribollet et al. 2006; Fricke et al. 2011; Charpy et al. 2012). Bruggemann et al. (1994c) reported cyanobacteria as a dominant component of the turf assemblages on the reef slopes of Bonaire, where we conducted our foraging observations. These epilithic and endolithic cyanobacteria are the primary nutritional resources for parrotfishes and are present in the majority of benthic samples targeted by many Indo-Pacific parrotfishes (Clements et al. 2016; Nicholson and Clements 2020). Excavating parrotfishes (e.g., *Sp. viride*)are able to access endolithic microalgae and cyanobacteria because their jaw morphology enables them to excavate reef substrate while grazing (Bellwood and Choat 1990; Bonaldo et al. 2014). The ability to excavate reef substrate also increases with body size in parrotfishes, meaning that large scrapers like *Sc. vetula* also have access to protein rich endoliths (Bellwood and Choat 1990; Bruggemann et al. 1994b; Adam et al. 2018). So, while we were unable to partition dietary targets within the algal turf communities from our video observations, it is likely that the majority of these bites also contained substantial cyanobacterial biomass.

In the Caribbean, *Sparisoma* spp. have often been described as macroalgal browsers (Adam et al. 2015b), and *Sp. aurofrenatum* is known to preferentially consume macroalgae, primarily *Dictyota* spp. (Dell et al. 2020). Consistent with these studies, we found that *Sp. viride* and *Sp. aurofrenatum* preferentially targeted macroalgae, with the majority of bites taken on *Dictyota* spp. (Fig. S3). It is possible that *Dictyota* spp. are targeted for their unusually high lipid content compared with other macroalgae (McDermid and Stuercke 2003). However, parrotfishes in our study were often observed spitting *Dictyota* spp. out after removing them from the substrate, a process that is important for the fragmentation and proliferation of *Dictyota* spp. at small spatial scales (Herren et al. 2006). Alternatively, parrotfishes could be attempting to remove epiphytic cyanobacteria that often grow upon fleshy macroalgae (Capone et al. 1977; Ballantine 1979; Gauna et al. 2015). This hypothesis remains untested, but warrants further investigation.

As cyanobacterial abundances increase on reefs globally (Ford et al. 2018), a knowledge gap has grown around trophic interactions involving BCMs, which are critical to consumer ecology and BCM dynamics. Our study provides further evidence that BCMs could be an important and preferred resource for parrotfishes. Intense grazing on BCMs by parrotfishes and other fishes may act as an important control on BCM proliferation. Future work should investigate variation in mat consumption (e.g., due to composition and palatability) and the effect of consumption on the trophic dynamics of BCMs.

## Acknowledgements

This research was supported by Florida State University (FSU) start-up funding and the Tatelbaum Ocean Research Fund awarded to S.J.M, as well as a Mote Research Assistantship from the William R. and Lenore Mote Eminent Scholar in Marine Biology Endowment at FSU and a Lerner-Gray Memorial Fund for Marine Research grant from the American Museum of Natural History awarded to J.C.M. We would like to thank C. Eckrich and R. Francisca for providing assistance in obtaining permits to work in the Bonaire National Marine Park. We would also like to thank the undergraduate field and lab assistants that assisted in data collection and processing: I. Basden, M. Dziewit, B. Clark, J. Portillo, B. Koechle, and J. Henson. Finally, we would like to thank K. Clements for valuable discussions regarding parrotfish nutritional ecology, and E. Cissell and the rest of the McCoy lab for feedback on earlier drafts of this manuscript.

