## Supplemental Information for "Preferential consumption of benthic cyanobacterial mats by Caribbean parrotfishes"

319 Stadium Drive

Tallahassee, FL 32306-4295

[2]University of North Carolina at Chapel Hill, Department of Biology

120 South Road

Chapel Hill, NC 27599-3280

Corresponding Author: Joshua C. Manning

ORCiD: 0000-0001-8130-5460

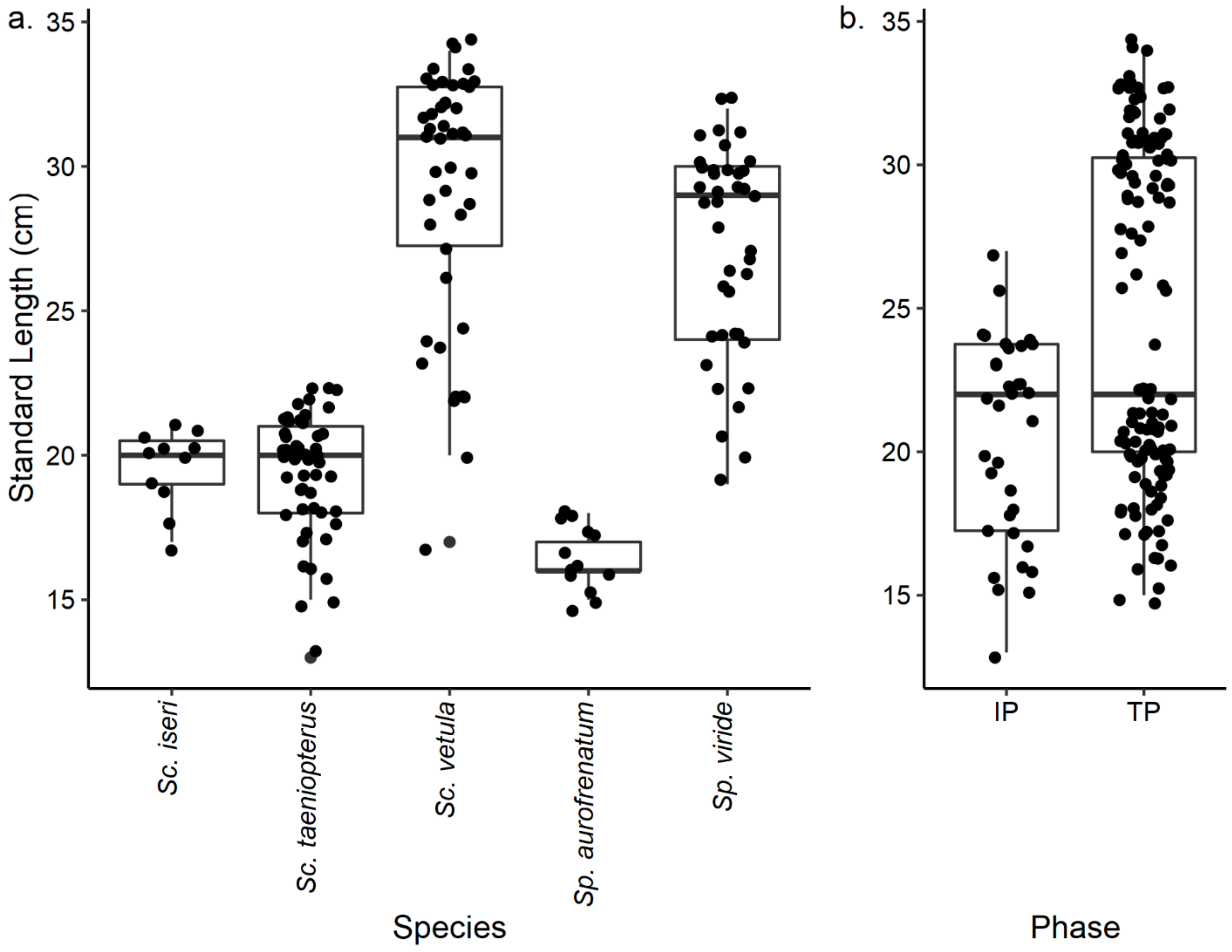


Fig. S1: Boxplots showing the median standard length (cm) of our five study species (a) and both ontogenetic phases (b). Dots represent individual fishes.

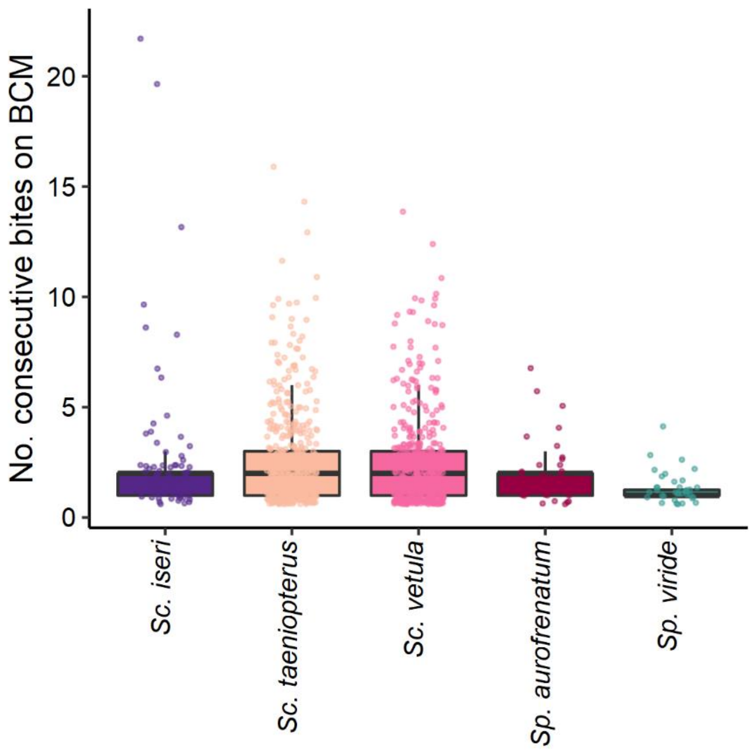


Fig. S2: A boxplot showing the median number of consecutive bites taken during feeding bouts on benthic cyanobacterial mats (BCM) by each of our five study species: *Scarus iseri*, *Scarus taeniopterus*, *Scarus vetula*, *Sparisoma aurofrenatum*, and *Sparisoma viride*. Dots represent individual feeding bouts.

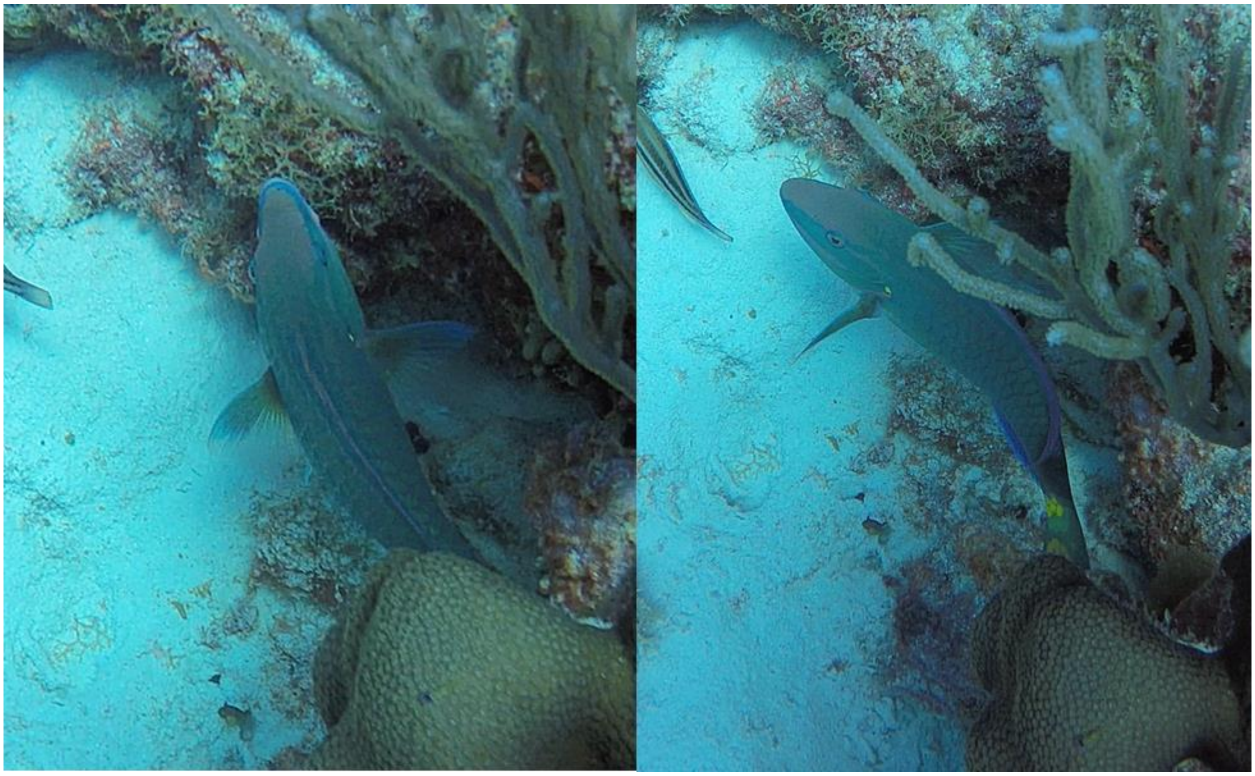

Fig. S3: A terminal phase *Sp. viride* removing *Dictyota* sp. from dead coral substrate.

Table S1: Mean ± SE percent cover of the benthic substrates at each study site estimated from 1/16 $m^2$ photoquadrats placed at 1 m intervals along 10 m transect lines that were placed haphazardly and run perpendicular to the reef slope at ~10 m depth.

| Site | N | Turfs & Crustose Coralline Algae | Live Coral | Sediment | Benthic Cyanobacterial Mats | Macroalgae |
|---|---|---|---|---|---|---|
| AC | 4 | 47.76 ± 4.23 | 22.31 ± 3.00 | 15.91 ± 5.78 | 8.56 ± 2.44 | 2.54 ± 0.43 |
| AQ | 5† | 63.59 ± 3.83 | 18.51 ± 5.22 | 14.38 ± 3.18 | 0.31 ± 0.20 | 0.57 ± 0.28 |
| BB | 4 | 47.60 ± 5.77 | 21.77 ± 6.17 | 21.75 ± 2.81 | 0.61 ± 0.30 | 0.16 ± 0.10 |
| IV | 4* | 38.66 ± 7.48 | 20.59 ± 5.29 | 36.12 ± 11.22 | 0.65 ± 0.13 | 0.30 ± 0.10 |
| TL | 4 | 49.01 ± 2.81 | 17.65 ± 5.93 | 15.97 ± 3.67 | 8.34 ± 2.75 | 2.97 ± 1.04 |
| **Site** | **N** | **Gorgonian** | **Sponge** | **Articulated Coralline Algae** | **Other Live** | **Unknown** |
| AC | 4 | 1.81 ± 0.71 | 0.36 ± 0.21 | 0.05 ± 0.05 | 0.11 ± 0.11 | 0.59 ± 0.29 |
| AQ | 5† | 1.28 ± 0.77 | 0.44 ± 0.27 | 0.18 ± 0.13 | 0.00 ± 0.00 | 0.75 ± 0.50 |
| BB | 4 | 6.21 ± 1.78 | 0.00 ± 0.00 | 0.05 ± 0.05 | 0.18 ± 0.18 | 1.66 ± 0.75 |
| IV | 4* | 2.54 ± 0.34 | 0.00 ± 0.00 | 0.11 ± 0.11 | 0.35 ± 0.18 | 0.67 ± 0.49 |
| TL | 4 | 4.67 ± 1.42 | 0.05 ± 0.05 | 0.15 ± 0.10 | 0.10 ± 0.06 | 1.09 ± 0.38 |

† At Aquarius (AQ), we included an additional unfinished transect (n=3 photoquadrats due to diving constraints); and another transect had only 9 photoquadrats.
*At Invisibles (IV), we inadvertently laid another photoquadrat at the 11 m mark on one transect, which we decided to include.

Table S2: Mean (± SE) density and biomass of parrotfishes estimated from eight 4 m x 25 m band transects at each study site: Angel City (AC), Aquarius (AQ), Bachelor's Beach (BB), Invisibles (IV), and The Lake (TL). Only estimates for terminal and initial phase parrotfishes are reported (fork length > 5 cm).

| **Site** | **Scientific Name** | **Density (# 100 m$^{-2}$)** | **Mass (g 100 m$^{-2}$)** |
|---|---|---|---|
| AC | *Scarus iseri* | 2.62 ± 1.07 | 65.01 ± 25.86 |
| | *Scarus taeniopterus* | 7.75 ± 0.82 | 871.27 ± 118.44 |
| | *Scarus vetula* | 1.25 ± 0.25 | 503.69 ± 163.13 |
| | *Sparisoma aurofrenatum* | 3.62 ± 0.53 | 190.27 ± 27.19 |
| | *Sparisoma viride* | 3.75 ± 0.73 | 456.66 ± 100.73 |
| AQ | *Scarus iseri* | 3.25 ± 0.80 | 85.35 ± 25.87 |
| | *Scarus taeniopterus* | 6.00 ± 1.16 | 615.19 ± 109.65 |
| | *Scarus vetula* | 2.00 ± 0.46 | 545.51 ± 130.34 |
| | *Sparisoma aurofrenatum* | 2.75 ± 0.59 | 124.68 ± 28.71 |
| | *Sparisoma chrysopterum* | 0.50 ± 0.50 | 22.93 ± 22.93 |
| | *Sparisoma viride* | 2.12 ± 0.83 | 470.00 ± 179.68 |
| BB | *Scarus iseri* | 2.50 ± 0.63 | 68.59 ± 15.96 |
| | *Scarus taeniopterus* | 6.88 ± 1.06 | 705.45 ± 84.37 |
| | *Scarus vetula* | 2.38 ± 0.46 | 800.74 ± 201.43 |
| | *Sparisoma aurofrenatum* | 2.62 ± 0.86 | 117.91 ± 40.84 |
| | *Sparisoma rubripinne* | 0.50 ± 0.38 | 100.69 ± 76.11 |
| | *Sparisoma viride* | 3.62 ± 0.50 | 801.83 ± 128.21 |
| IV | *Scarus iseri* | 3.38 ± 0.60 | 132.76 ± 23.43 |
| | *Scarus taeniopterus* | 6.88 ± 1.01 | 652.65 ± 118.51 |
| | *Scarus vetula* | 1.25 ± 0.73 | 427.10 ± 205.02 |
| | *Sparisoma aurofrenatum* | 4.00 ± 0.80 | 204.52 ± 33.08 |
| | *Sparisoma chrysopterum* | 0.12 ± 0.12 | 35.59 ± 35.59 |
| | *Sparisoma viride* | 2.25 ± 0.82 | 452.97 ± 173.26 |
| TL | *Scarus iseri* | 3.25 ± 0.96 | 119.35 ± 38.44 |
| | *Scarus taeniopterus* | 7.25 ± 1.28 | 751.92 ± 135.73 |
| | *Scarus vetula* | 2.00 ± 0.38 | 698.68 ± 167.63 |
| | *Sparisoma aurofrenatum* | 3.62 ± 0.75 | 172.08 ± 32.10 |
| | *Sparisoma viride* | 2.38 ± 0.46 | 652.90 ± 217.62 |

Table S3: Mean (± SE) standard length, total time followed (total time), time the fish were not clearly visible in the video (lost time), and observation time (total time – time lost) for terminal phase (TP) and initial phase (IP) parrotfishes of our five study species: *Scarus iseri*, *Scarus taeniopterus*, *Scarus vetula*, *Sparisoma aurofrenatum*, and *Sparisoma viride*.

| **Species** | **Phase** | **N** | **Standard Length (cm)** | **Total Time (min)** | **Time Lost (min)** | **Observation Time (min)** |
|---|---|---|---|---|---|---|
| *Sc. iseri* | TP | 11 | 19.6 ± 0.4 | 14.57 ± 0.47 | 0.03 ± 0.02 | 14.54 ± 0.48 |
| *Sc. taeniopterus* | IP | 11 | 16.4 ± 0.5 | 10.62 ± 0.29 | 0.09 ± 0.05 | 10.53 ± 0.32 |
| | TP | 41 | 20.1 ± 0.2 | 10.47 ± 0.17 | 0.18 ± 0.05 | 10.29 ± 0.17 |
| *Sc. vetula* | IP | 11 | 22.4 ± 0.7 | 11.04 ± 0.18 | 0.03 ± 0.02 | 11.01 ± 0.18 |
| | TP | 35 | 31.4 ± 0.3 | 15.42 ± 0.24 | 0.03 ± 0.03 | 15.39 ± 0.23 |
| *Sp. aurofrenatum* | TP | 13 | 16.5 ± 0.3 | 13.35 ± 0.42 | 0.13 ± 0.05 | 13.22 ± 0.41 |
| *Sp. viride* | IP | 12 | 22.7 ± 0.6 | 11.15 ± 0.34 | 0.11 ± 0.05 | 11.04 ± 0.34 |
| | TP | 28 | 29.1 ± 0.4 | 15.59 ± 0.24 | 0.08 ± 0.03 | 15.51 ± 0.25 |
